# Structural basis for context-specific inhibition of translation by oxazolidinone antibiotics

**DOI:** 10.1101/2021.08.10.455846

**Authors:** Kaitlyn Tsai, Vanja Stojković, D. John Lee, Iris D. Young, Teresa Szal, Nora Vazquez-Laslop, Alexander S. Mankin, James S. Fraser, Danica Galonić Fujimori

**Affiliations:** Department of Cellular and Molecular Pharmacology; University of California San Francisco, San Francisco, CA 94158, USA; Department of Bioengineering and Therapeutic Sciences; University of California San Francisco, San Francisco, CA 94158, USA; Department of Pharmaceutical Sciences, University of Illinois at Chicago, Chicago, IL 60607, USA; Center for Biomolecular Sciences, University of Illinois at Chicago, Chicago, IL 60607, USA; Quantitative Biosciences Institute, University of California San Francisco, San Francisco, CA 94158, USA; Department of Pharmaceutical Chemistry, University of California San Francisco; San Francisco, CA 94158, USA

**Author notes:** Authors contributed equally to this work (Kaitlyn Tsai, Vanja Stojković, D. John Lee).

**Keywords:** linezolid, radezolid, oxazolidinone antibiotics, peptidyl transferase center, cryo-EM, stalled ribosome, nascent peptide, selective inhibition of translation, Cfr, m^8^A2503

## Abstract

The antibiotic linezolid, the first clinically approved member of the oxazolidinone class, inhibits translation of bacterial ribosomes by binding to the peptidyl transferase center. Recent work has demonstrated that linezolid does not inhibit peptide bond formation at all sequences but rather acts in a context-specific manner, namely when alanine occupies the penultimate position of the nascent chain. In this study, we determined that the second-generation oxazolidinone radezolid also induces stalling with alanine at the penultimate position. However, the molecular basis for context-specificity of these inhibitors has not been elucidated. In this study, we determined high-resolution cryo-EM structures of both linezolid and radezolid-stalled ribosome complexes. These structures reveal that the alanine side chain fits within a small hydrophobic crevice created by oxazolidinone, resulting in improved ribosome binding. Modification of the ribosome by the antibiotic resistance enzyme Cfr disrupts stalling by forcing the antibiotic to adopt a conformation that narrows the hydrophobic alanine pocket. Together, the structural and biochemical findings presented in this work provide molecular understanding of context-specific inhibition of translation by clinically important oxazolidinone antibiotics.

## INTRODUCTION

During translation, peptide bond formation occurs within the peptidyl transferase center (PTC) of the ribosome. The PTC is located within the large ribosomal subunit and catalyzes extension of the polypeptide chain through proper positioning of the peptidyl-tRNA in the P-site and aminoacyl tRNA in the A-site. Due to its functional importance, the PTC of the bacterial ribosome is a common target for antibiotics that inhibit translation^1,2^.

The PTC antibiotic linezolid (LZD, **Fig. 1a**) was the first clinically approved member of the synthetic oxazolidinone class of antibiotics^3^. Linezolid is used to treat drug-resistant gram-positive infections including those caused by methicillin-resistant *S. aureus* and vancomycin-resistant Enterococci^4^. Initial cross-linking experiments identified the binding site for linezolid within the PTC A-site only when performed with actively translating ribosomes^5,6^, suggesting that other translation components may be involved in linezolid binding. However, existing structures of linezolid-bound ribosomes have been obtained with the ribosomes devoid of charged tRNAs^7,8^ or ribosomes containing only a P-site tRNA lacking a nascent chain^9^. Recent evidence obtained through ribosome profiling and single-molecule studies demonstrated that linezolid does not indiscriminately inhibit the formation of every peptide bond but rather interferes with translation at certain mRNA sites. Robust inhibition of translation and ribosome stalling by linezolid is strongly favored when the amino acid alanine occupies the penultimate, (−1), position within the nascent chain^10,11^. Together, these results suggest that interactions between linezolid and the nascent peptide may be important for stabilizing antibiotic binding to the ribosome. However, the exact nature of these interactions is yet to be elucidated.

**Figure 1.**
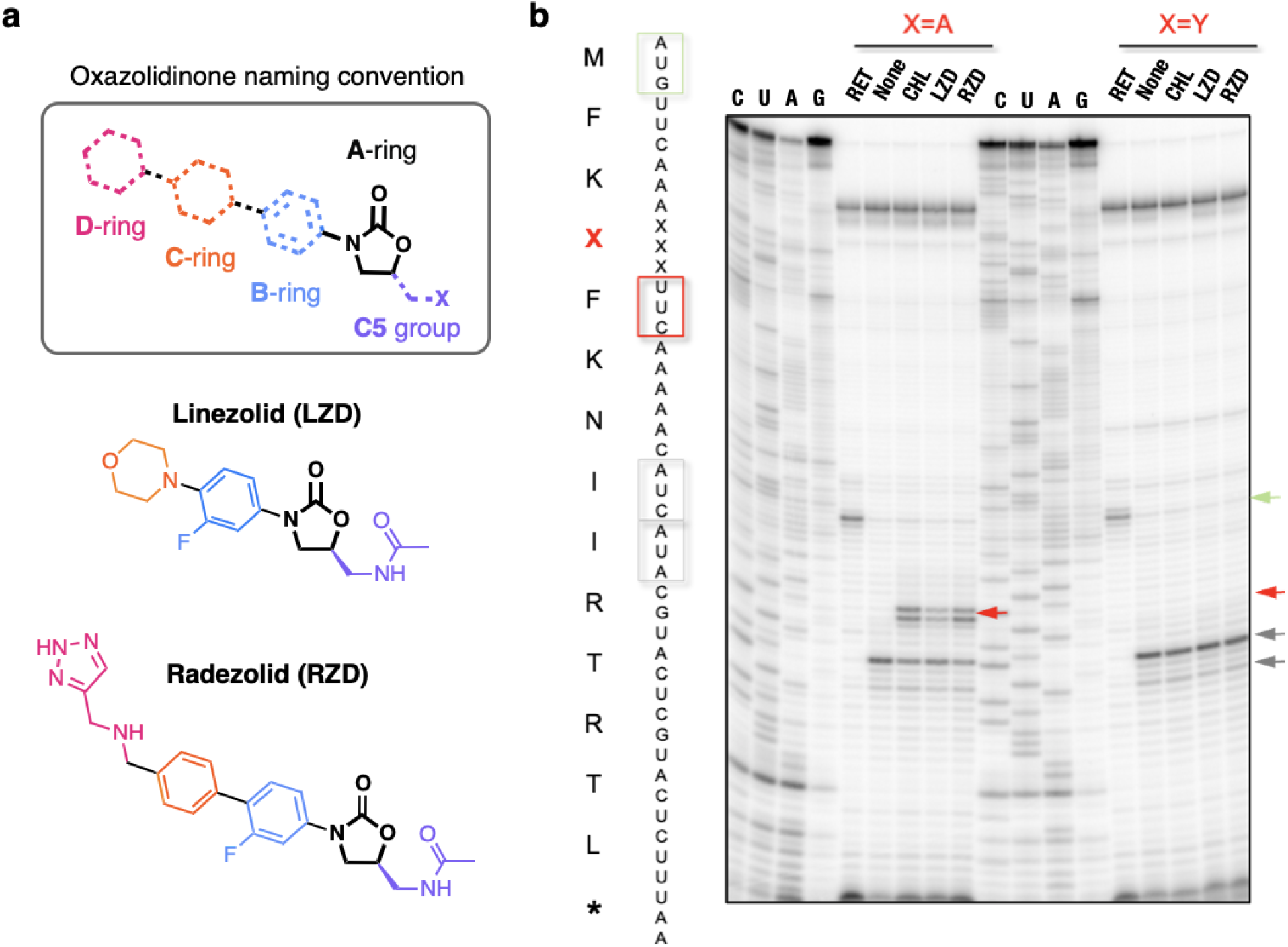
Radezolid induces ribosome stalling with alanine in the penultimate position. (**a**) Molecular architecture of oxazolidinone antibiotics. The oxazolidinone (A-ring) portion of the molecule is conserved amongst the oxazolidinone class and designated in black. Chemical moieties that vary amongst oxazolidinone derivatives, the C5-moiety, and B, C, and D rings, are designated by their respective color. (**b**) Toeprinting assays performed on model mRNAs encoding stalling and non-stalling peptides. The control antibiotic retapamulin (RET) was used to stall ribosomes at the start codon indicated by the green arrow^15^. ‘None’ designates samples lacking ribosome-targeting antibiotics. The control antibiotic chloramphenicol (CHL) was used to stall with alanine at the penultimate position^10,11^. The toeprint bands corresponding to the prominent stall sites observed in reactions containing chloramphenicol (CHL), LZD, or RZD when the 4th amino acid is an alanine, but not tyrosine, are indicated by the red arrow. Due to the inclusion of the Ile-tRNA synthetase inhibitor mupirocin in all toeprinting reactions, any ribosomes not stalled at an upstream codon are trapped at the downstream Ile codons designated by grey arrows.

The clinical success of LZD and emergence of resistance to this antibiotic through modification of the ribosome by chloramphenicol-florfenicol resistance enzyme Cfr sparked the development of second-generation derivatives, such as radezolid (RZD), with improved potency^12^. Radezolid (**Fig. 1a**) is in clinical development for bacterial acne and community-acquired pneumonia^13^. Compared to linezolid, radezolid retains the aryl-oxazolidinone core (rings A and B) and C5 group but has alterations to the C/D ring system. Given that key chemical elements are conserved between LZD and RZD it is plausible that RZD can also act as a context-specific inhibitor of translation. The importance of both conserved and distinct structural features of second generation oxazolidinones to translation inhibition and context specificity, if any, is yet to be elucidated.

In this work, we discovered that RZD exhibits ribosome stalling behavior similar to that of LZD, arresting translating ribosomes when alanine occupies the penultimate position within the nascent peptide chain. Capitalizing on the stalling preference of these two antibiotics, we generated high resolution cryo-EM structures of LZD- and RZD-stalled ribosome complexes. Direct comparison of the drug-stalled translation complexes with the structures of the vacant ribosomes bound to the same antibiotics enabled identification of molecular contacts that improve oxazolidinone binding to the ribosome. Specifically, we find that the penultimate alanine fits snugly within a shallow hydrophobic pocket created by the oxazolidinone molecules, providing structural rationale for context specificity. Our analysis of RZD action on ribosomes modified by the oxazolidinone resistance enzyme Cfr and the structure of RZD bound to the Cfr-modified ribosome provides the first insights into how this second generation oxazolidinone interacts with a LZD-resistant ribosome.

## RESULTS

### Linezolid and radezolid have similar ribosome stalling behavior

Previous *in vitro* toe-printing experiments have demonstrated that LZD induces ribosome stalling on a model mRNA sequence when the amino acid alanine is located in the penultimate (−1) position of the nascent polypeptide chain^10,11^ (**Fig. 1b**). LZD-induced ribosome stalling is abolished when the penultimate alanine is replaced with tyrosine (**Fig. 1b**). To evaluate if the second generation oxazolidinone RZD shows similar stalling behavior, we carried out *in vitro* toeprinting analysis to monitor the position of stalled ribosomes on the previously described mRNAs encoding the following peptide sequences: MFK**A**FKNIIRTRTL and MFK**Y**FKNIIRTRTL^11^.

Similarly to LZD, the presence of RZD permits formation of the first peptide bond with both mRNA transcripts as templates (**Fig. 1b**). This result indicates that RZD is not a universal inhibitor of translation and also does not act as initiation inhibitor as suggested for LZD by some earlier studies^14^. While no inhibition of translation at the early mRNA codons was observed on MFK**Y**FK-encoding template, presence of RZD or LZD led to selective stalling of the ribosome during the translation of MFK**A**FK-encoding template. The stalling occurred at the F5 codon when an alanine residue appeared in the penultimate position within the nascent peptide chain. These results indicate that RZD- or LZD-bound ribosomes are unable to catalyze peptide bond formation between F5 and K6 when the MFK**A**F nascent peptide occupies the exit tunnel. Comparison of the relative intensity of the stalled ribosome toeprint bands suggests that RZD is a stronger inducer of ribosome stalling than LZD. The observation that RZD is unable to stall the ribosome if the critical alanine residue is replaced with a tyrosine (MFK**Y**F) is consistent with the specificity of action of LZD (**Fig. 1b**).

### Generation of stalled ribosomes complexes and cryo-EM analysis

Guided by the *in vitro* stalling behavior, we designed a stalling peptide to capture LZD and RZD stalled ribosome complexes (SRC) for structural analysis, herein referred to as LZD-SRC and RZD-SRC, respectively. Stalled complexes were generated by conducting coupled *in vitro* transcription-translation reactions with *E. coli* ribosomes in the presence of oxazolidinone antibiotics (**Fig. 2a**). To bias formation of ribosomes with nascent peptide stalled at the F5 codon, we designed the DNA template where the open reading frame encoding the MFK**A**F stalling peptide lacked a stop codon (**Fig. 2a, Supplementary Fig. 1**). Stalled 70S ribosome complexes were purified away from other components of the translation reaction by sucrose gradient fractionation and vitrified on carbon grids for cryo-EM analysis. Refinement and reconstruction was performed using the cisTEM software suite^16^ to obtain 2.5 Å resolution structure of LZD-SRC and 2.5 Å resolution of RZD-SRC (**Fig. 2b, Supplementary Fig. 2, Supplementary Table 1**).

**Figure 2.**
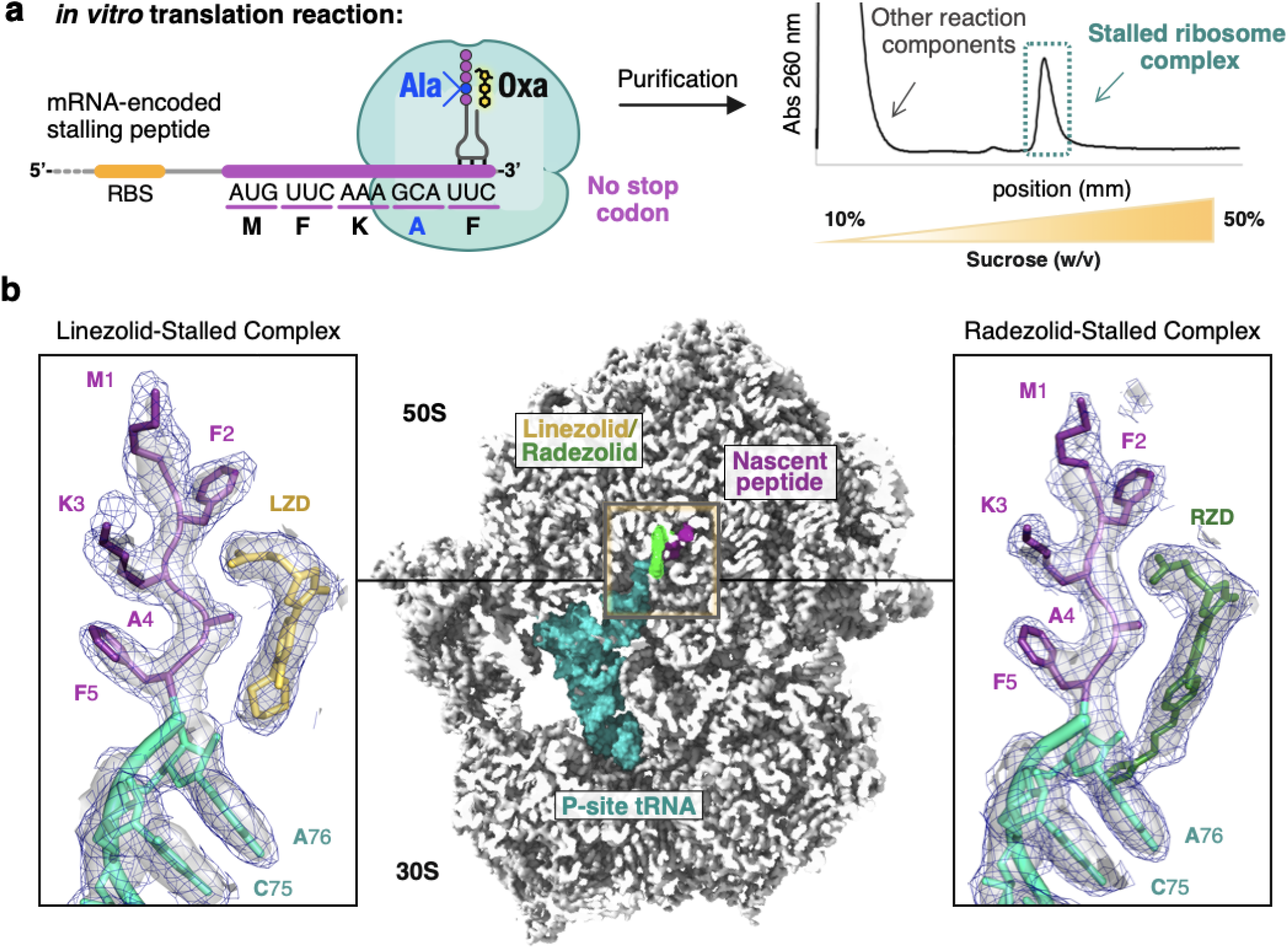
Cryo-EM structures of linezolid and radezolid-stalled ribosome complexes. (**a**) Stalled complexes were generated by performing coupled *in vitro* transcription-translation reactions in the presence of the oxazolidinone (Oxa) antibiotic linezolid or radezolid. Complexes were further purified by sucrose gradient fractionation. (**b**) Cross-section of the cryo-EM density map of the 70S ribosome in complex with peptidyl-tRNA and oxazolidinone. Inserts are close-up views of the linezolid (LZD) or radezolid (RZD) in complex with the MFK**A**F nascent peptide. Coulomb potential density is contoured at 4.0**σ** in surface representation and 1.0**σ** in mesh representation. Figure was made using unsharpened maps.

The generated cryo-EM maps of LZD-SRC and RZD-SRC have well-defined densities for the oxazolidinone antibiotic and rRNA nucleotides, especially those within the PTC. Both cryo-EM maps also have well defined densities for the peptidyl-tRNA located in the P-site. No A-site or E-site tRNAs are present. Modeling of the density maps unambiguously assigns the penultimate residue in the nascent peptide as alanine, as expected based on the DNA template used in the experiment (**Fig. 2b, Supplementary Fig. 2**). This assignment is further supported by cryo-EM densities corresponding to mRNA:tRNA interaction. The codon-anticodon densities are best modeled by the interaction between UUC:GAA-tRNA^Phe^ rather than GCA:UGC-tRNA^Ala^ (**Supplementary Fig. 3**).

To directly compare how presence of the nascent peptide may influence positioning of the antibiotic and/or conformation of rRNA nucleotides, we also generated structures of vacant *E. coli* 50S ribosomal subunits in complex with LZD or RZD alone, herein referred to as LZD-50S and RZD-50S, at 2.4 Å resolution and 2.5 Å resolution, respectively (**Supplementary Fig. 2, Supplementary Table 1**). Similarly to the stalled complexes, antibiotic-only bound structures have unambiguous densities for the oxazolidinone antibiotic.

### The penultimate alanine facilitates oxazolidinone antibiotic binding to the ribosome

The overall binding modes of LZD and RZD within the PTC in antibiotic-only bound and stalled ribosome complexes are similar to those described previously^7–9,17^. The fluorophenyl moiety (B-ring) sits in the A-site cleft, a hydrophobic pocket formed by splayed out nitrogen bases of nucleotides C2452 and A2451 (**Supplementary Fig. 4a**,**b**). The oxazolidinone ring (A-ring) of LZD and RZD is positioned in an offset π-π stacking interaction with Ψ2504. Interestingly, we find that the carbonyl of the oxazolidinone ring does not interact with rRNA but is rather chelated to what is likely a Mg^2+^ ion based on the coordination geometry (**Supplementary Fig. 4c**,**d**), corroborating previous SAR studies that demonstrated the importance of an electron-pair donor for activity^18^.

The binding poses for both LZD and RZD are near-identical between the antibiotic-only and stalled-ribosome structures (**Supplementary Fig. 5**). We do, however, observe improved density for both antibiotics, suggesting that presence of the stalling nascent peptide stabilizes the placement of LZD and RZD in the ribosome (**Fig. 3a,b**). Specifically, we observe improved density for the C5 acetamide group for both antibiotics, as well as enhanced density for the D-ring of RZD in the stalled structures. Of note, in previously published density maps^7–9,17^, the acetamide is less well resolved and has been modeled in a variety of positions, likely due to its ability to sample multiple conformations. Our structures suggest that the nascent peptide present in the stalled ribosome complex stabilizes the C5 group thereby providing a more biologically relevant view on the drug placement.

**Figure 3.**
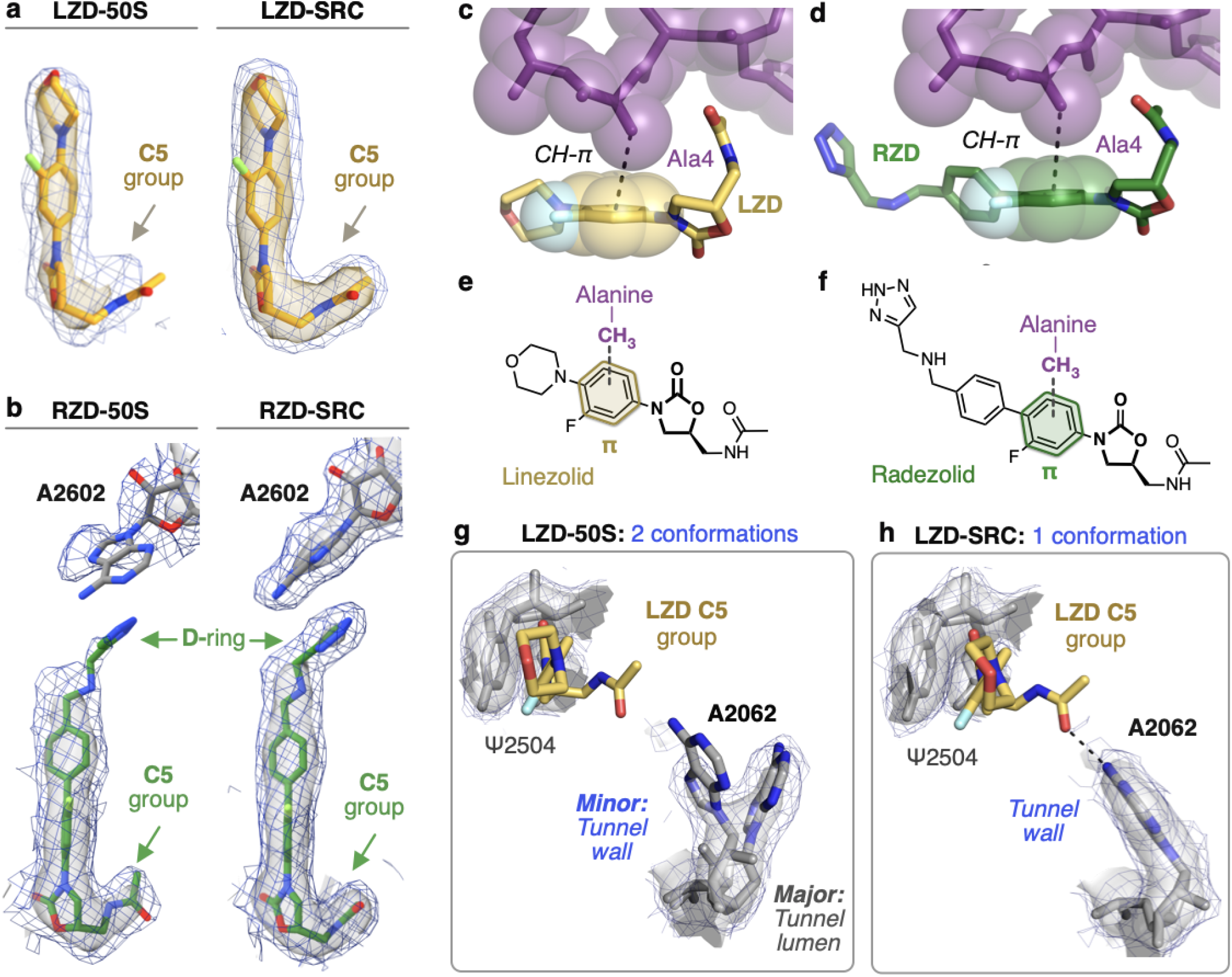
Stalling peptide stabilizes oxazolidinone binding to the ribosome. For this figure, coulomb potential density is contoured at 4.0σ in surface representation and 1.0σ in mesh representation from unsharpened maps generated with a soft 50S mask. (**a**) Comparison of linezolid density in the linezolid-only bound (LZD-50S) and the linezolid-stalled complex (LZD-SRC). (**b**) Comparison of radezolid density in the radezolid-only bound (RZD-50S) and the radezolid-stalled complex (RZD-SRC). (**c**) Close up view of the CH-π interaction between the B-ring of LZD and the penultimate alanine (Ala4). (**d**) Close up view of the CH-π interaction between the RZD B-ring and the penultimate alanine. (**e**,**f**) Schematic of the CH-π interaction involved in stabilizing antibiotic binding in (**e**) LZD-stalled and (**f**) RZD-stalled ribosome complexes. (**g**) Density of the exit tunnel rRNA nucleotide A2062 in LZD-only bound structure, highlighting two conformations. (**h**) Density of A2062 in the LZD-stalled structure, with only one observed conformation.

The nascent protein chain, and more specifically, the penultimate Ala residue, contributes directly to formation of the drug binding site (**Fig. 2b, Fig. 3c,d**). In contrast to the orientation of the flanking residues within the nascent peptide, the side chain of the penultimate alanine faces towards the oxazolidinone binding pocket. In this orientation, the methyl group of alanine fits snugly within the hydrophobic crevice formed by the C5 group and A/B-ring system of the antibiotic. The crevice is deep enough to accommodate the small side chain of alanine, but is too shallow to fit bulkier side chains, including that of tyrosine, providing rationale for differential stalling preferences observed in our toeprinting experiments (**Fig. 1b**). In the stalled complex, the alanine’s side chain methyl group engages in a CH-π interaction with the aryl B-ring of the oxazolidinone (3.6 Å and 3.9 Å between C atom and plane of the B-ring for LZD-SRC and RZD-SRC, respectively) (**Fig. 3c,d**). This interaction likely facilitates antibiotic binding to the A-site of the PTC, resulting in enhanced occlusion of incoming aminoacyl tRNAs. Glycine, the smallest amino acid residue and which lacks a side chain, does not clash with the ribosome-bound antibiotic when present in the penultimate position of the nascent peptide, yet it is not favored for drug-induced ribosome stalling^10,11^. The inability of glycine to form the alanine-specific CH-π interaction with the drug molecule likely explains this observation.

A similar CH-π interaction has been recently observed between the penultimate alanine and aryl ring of chloramphenicol (CHL), another PTC-targeting antibiotic that also exhibits context-specific inhibition of translation^19^. In contrast to CHL, which also induces robust ribosome stalling when serine or threonine occupy the penultimate position, LZD exhibits a strong preference for alanine^10^. While *in silico* modeling of serine or threonine at the penultimate position in LZD-SRC reveals that serine can be accommodated, the methyl group of threonine generates a steric clash for all favored rotamers (**Supplementary Fig. 6**). Recent work hypothesized that CHL-induced stalling with serine in the penultimate position of the nascent peptide is stabilized by a H-bond between the serine hydroxyl and chlorine atom of CHL^19^. Since the C5 group of LZD does not have an analogous electron-pair donor, it is likely that the unsatisfied H-bond acceptor, rather than sterics, accounts for LZD’s strong preference for alanine over serine.

### Dynamic nucleotides are stabilized to provide additional interactions with the oxazolidinone antibiotics

In LZD and RZD antibiotic-only 50S structures, the exit tunnel nucleotide A2062 can adopt two distinct conformations (**Fig. 3g**). In its dominant conformation the A2062 base projects into the lumen of the exit tunnel, while in the minor conformation, A2062 is rotated and lays flat against the tunnel wall. Strikingly, in the stalled ribosome complexes with the nascent protein chain occupying the tunnel, the A2062 base is stabilized in the rotated state juxtaposed against the tunnel wall (**Fig. 3h, Supplementary Fig. 7a**). The rotated state of A2062 is stabilized by a H-bond between the N1 atom of adenine and the stalling peptide backbone, as well as a non-canonical A:A base-pair with m^2^A2503 (**Supplementary Fig. 7b**). In this conformation, the exocyclic amine of A2062 is within H-bonding distance of the acetamide carbonyl of LZD/RZD (2.7 Å/3.3 Å), which likely explains why we observe improved density for the C5 group in the stalled complexes (**Fig. 3a,b**). This interaction has not been observed in existing structures of oxazolidinone-bound vacant ribosomes due to the alternative orientation of A2062. The peptide-induced interaction between A2062 and the oxazolidinone is distinct from that observed with CHL. In structures of chloramphenicol without nascent peptide, A2062 is already in a rotated state to form a H-bond with the antibiotic^20^.

Compared to the antibiotic-only bound structures, we also observe stabilization of dynamic PTC nucleotides in conformations that provide additional contacts with the antibiotics. Notably, the global resolutions of the maps are similar and many residues show nearly identical fits to density at the same map threshold (**Supplementary Fig. 8**). We observe improved density for U2585, which as the C4 enol tautomer could provide a H-bonding interaction with the oxygen atom of the morpholine ring in LZD (**Supplementary Fig. 7, 8**). Although U2506 does not make direct contact with the LZD, we observe dramatically improved density for this nucleotide in a conformation analogous to an uninduced or nonproductive state of the PTC^21^ (**Supplementary Fig. 7c, 8**). In the RZD-stalled complex, we observe improved densities for U2506, U2585 and A2602 (**Supplementary Fig. 8**). Nucleotides U2506 and A2602 provide π-π stacking interactions with the C- and D-ring, respectively, while nucleotide U2585 engages in a H-bond with the secondary amine of RZD (**Fig. 3b, Supplementary Fig. 7**). Interestingly, the D-ring interaction with A2602 has not been observed in a previous ribosome structure with RZD ^17^ but likely explains why RZD is a better inhibitor of translation compared to LZD, as the stabilized D-ring would provide additional steric interference with aminoacyl tRNA binding (**Supplementary Fig. 9**). Together these results suggest that, in addition to favorable interactions with the penultimate alanine, interactions between rRNA nucleotides and the oxazolidinone likely play a role in improving antibiotic binding to the ribosome to facilitate stalling.

### Cfr methylation destabilizes the stalled ribosome complex

A prevalent resistance mechanism to LZD identified in multiple clinical isolates worldwide^22–28^ involves methylation of rRNA by the Cfr enzyme, which adds a methyl group at the C8 atom of A2503 (m^8^A2503) in 23S rRNA^29–33^. The Cfr modification disrupts LZD binding to the ribosome by introducing a steric clash between the installed methyl mark and the C5 group of the antibiotic. While this modification confers high levels of resistance to LZD, RZD retains modest efficacy against Cfr-positive strains^34,35^, likely due to retained interactions on the other side of the molecule involving the D-ring (**Fig. 3b**). However, this suggests that when RZD binds to Cfr-modified ribosomes, the C5 group must adopt an alternative conformation to accommodate C8 methyl group of m^8^A2503. Given that the C5 group is an important component of the antibiotic pocket required to fit the alanine side chain, we wanted to evaluate the ability of RZD to induce stalling of Cfr-modified ribosomes.

To perform *in vitro* assays, we expressed Cfr in *E. coli* and isolated ribosomes with near-complete methylation of m^8^A2503 as described previously^33^. As expected, RZD retains activity against the m^8^A2503 ribosomes. *In vitro* translation of the sf-GFP by Cfr-modified ribosomes could not be suppressed by LZD but was readily inhibited by RZD (IC_50_ ∼1 µM (**Supplementary Fig. 10a**)). As expected, in toe-printing experiments the m^8^A2503 modification alleviated LZD-induced ribosome arrest at the F5 codon of the MFK**A**F…-encoding ORF (**Fig. 4a**). Strikingly, RZD, while retaining its general translation inhibitory activity, also failed to arrest the m^8^A2503 ribosome at the F5 codon of the MFK**A**F template (**Fig. 4a**). Together, these results suggest that the presence of C8 methyl group at A2503 does not prevent antibiotic binding but alters its ability to arrest translation in the penultimate alanine-specific manner.

**Figure 4.**
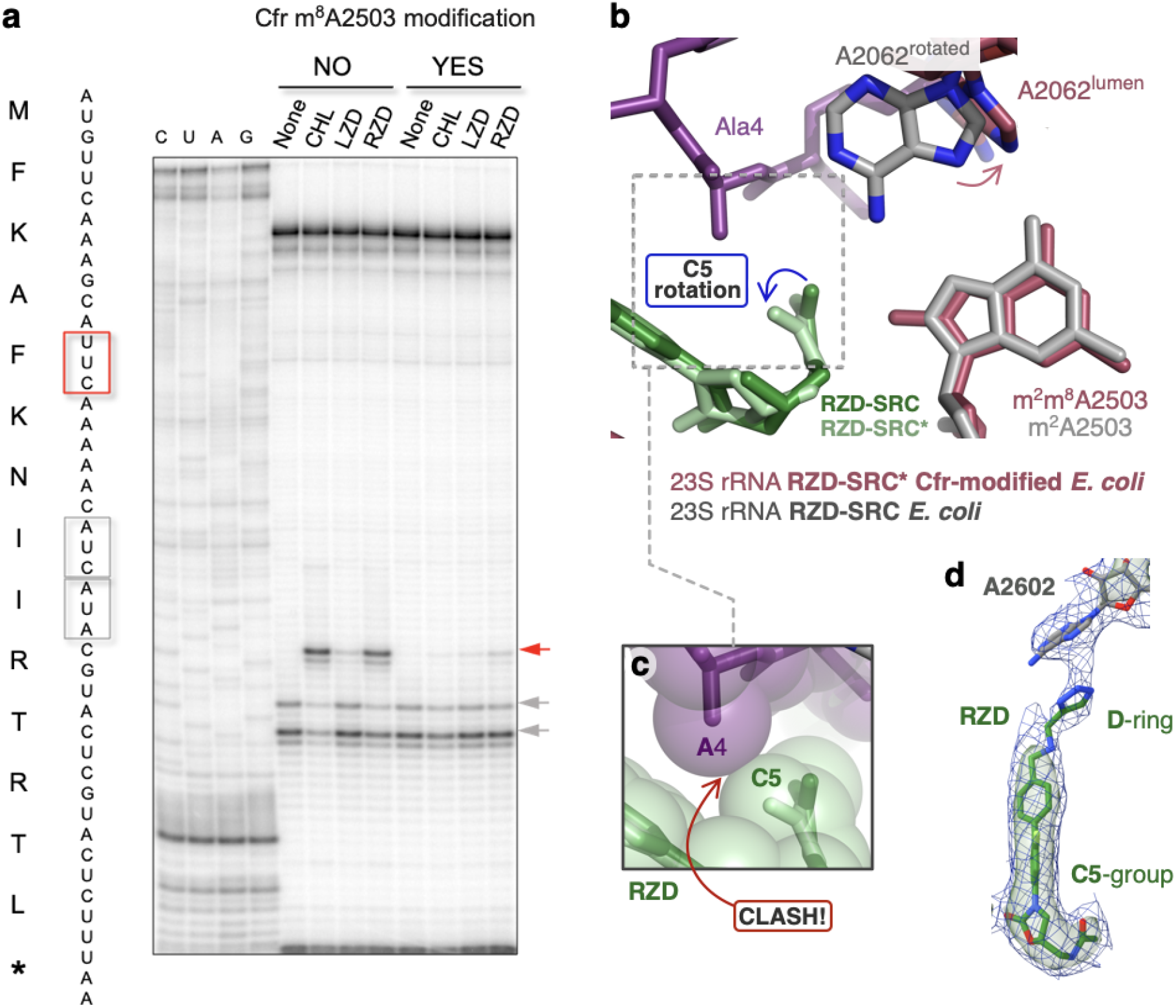
The Cfr modification m^8^A2503 perturbs linezolid and radezolid-induced ribosome stalling. (**a**) Toeprinting assays performed on the MFK**A**F stalling peptide sequence. Prominent stall sites observed in reactions containing chloramphenicol (CHL), linezolid (LZD), or radezolid (RZD) with *wildtype* ribosomes, but not with Cfr-modified ribosomes, are indicated by the red arrow. Due to the inclusion of the tRNA synthetase inhibitor mupirocin in all toeprinting reactions, any ribosomes not stalled at an upstream codon are forced to stall at the Ile codons designated by grey arrows. (**b**) Structural rearrangements identified in RZD-forced stalled complex with Cfr-modified ribosomes (RZD-SRC*). Stalling peptide in purple is from RZD-SRC. Rotation of the RZD C5 group is required to accommodate m^8^A2503, resulting in a steric clash between the penultimate alanine and the C5 group, close-up view shown in sphere representation in panel (**c**). (**d**) Retained interaction between RZD D-ring and A2602 in Cfr-modified ribosomes.

In search of a molecular explanation for this result, we determined the structure of RZD-bound to Cfr-modified ribosome ‘forced-stalled’ with the MFK**A**F-tRNA in the P-site at 2.4 Å resolution. While the overall binding of the antibiotic molecule was similar to that observed in the stalled *wildtype* ribosome, the C5 group of RZD bound to the m^8^A2503 ribosome adopts a conformation distinct from that of the *wildtype* complex (**Fig. 4b**). In this alternative conformation, the C5 group encroaches upon the binding pocket of the penultimate alanine to introduce a mild steric clash (**Fig. 4c**). As a result, the oxazolidinone cavity becomes a poor fit for the penultimate alanine of the nascent protein chain. Destabilization of a properly stalled complex is exhibited by poor density for certain side chains of the nascent peptide (**Supplementary Fig. 10b**). We also observe reversion of exit tunnel nucleotide A2062 to its lumen conformation which is now positioned too far away to engage in a H-bond with RZD. These results suggest that the m^8^A2503 modification alters the RZD stalling landscape by changing the interface between the antibiotic and the nascent peptide.

Our structural findings also provide important insights into how RZD retains efficacy against LZD-resistant ribosomes. While the Cfr modification perturbs positioning of the C5 group, which likely disrupts interaction with exit tunnel nucleotide A2062, we observe retention of other interactions. Although somewhat diminished, we observe densities for the D-ring and A2602 engaged in a π-π stacking interaction analogous to that observed in the *wildtype*, non-Cfr modified complex (**Fig. 4d**). LZD does not contain the additional D-ring and is thus more dramatically impacted by lost interactions due to m^8^A2503. Our results suggest that in addition to shortening of the C5 group, which has been carried out with other oxazolidinones derivatives^13,36^, extension of the ring system on the opposite end of the molecule is an orthogonal, viable strategy for generating oxazolidinone antibiotics that overcome Cfr resistance.

## DISCUSSION

We identified two factors that contribute to context specificity of oxazolidinones LZD and RZD (**Fig. 5**). While the antibiotic can still bind to the ribosome that lacks the nascent protein chain, as revealed by our and previously published structures of the vacant ribosome complexed with LZD and RZD, the presence of the nascent chain stabilizes binding of the drug by providing additional points of contact. Most importantly, the side chain of the penultimate alanine residue is intercalated into the complementary-shaped cavity formed by the drug molecule (**Fig. 2b, Fig. 3c,d**). Larger amino acids in the penultimate position of the nascent peptide would clash with the antibiotic, preventing its binding. In contrast, glycine cannot make a CH-π interaction with the drug, thereby making its binding less favorable. As a secondary effect of the alanine interaction, we also observe stabilization of dynamic nucleotides in conformations that provide additional contacts with the antibiotic (**Supplementary Fig. 7**). The culmination of these interactions leads to improved antibiotic binding to the ribosomal A-site in the presence of nascent peptides containing a penultimate Ala, facilitating competition with incoming aminoacyl tRNAs and thus resulting in ribosome stalling.

**Figure 5.**
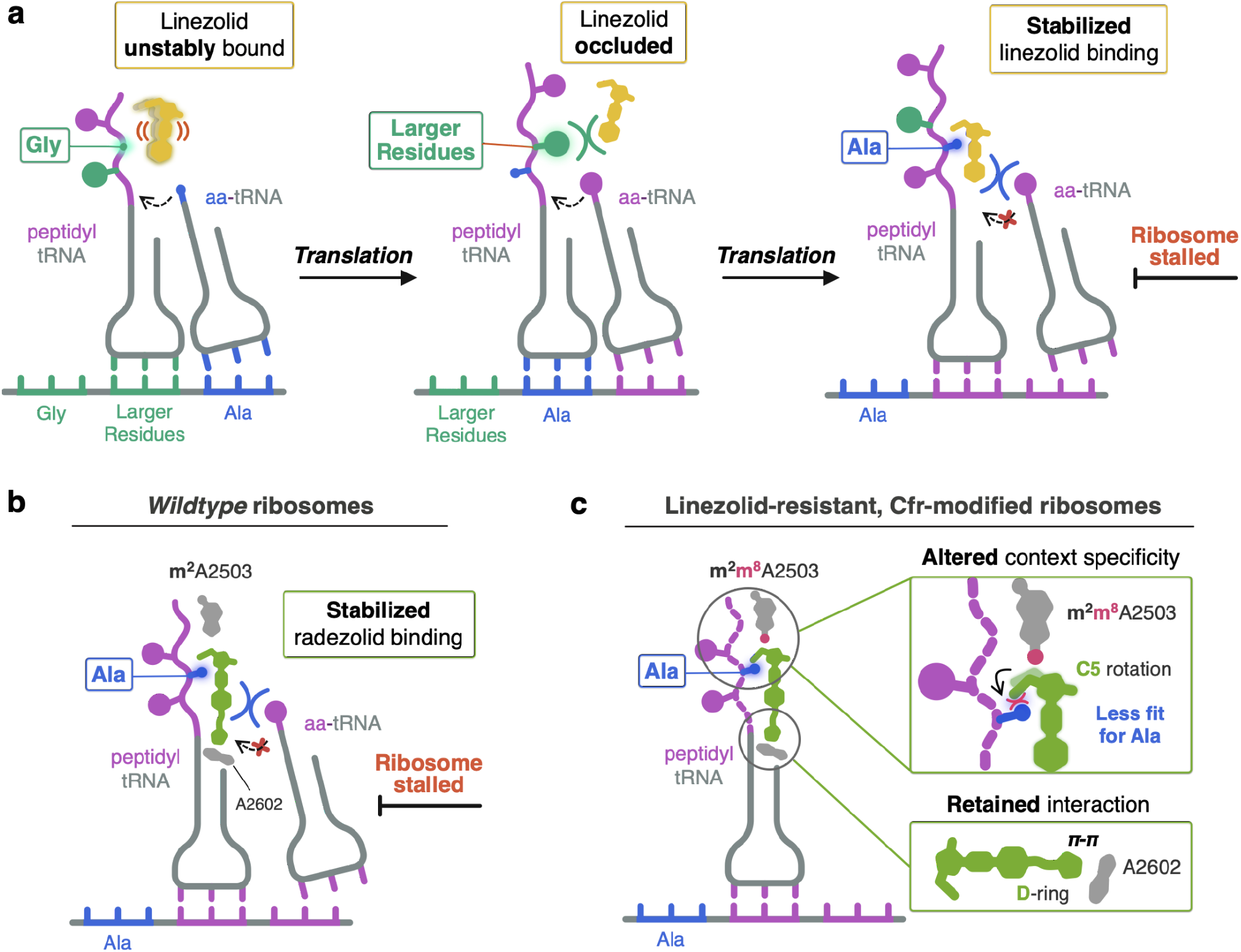
Model for oxazolidinone context-specific inhibition of translation. (**a**) Linezolid (yellow) is either unstably bound or sterically occluded from binding to the ribosome when glycine (Gly) or larger residues occupy the penultimate position within the nascent chain. When alanine (Ala, blue) occupies the penultimate position, linezolid becomes stably bound to the ribosomal PTC A-site, enabling linezolid to better compete away incoming aminoacyl tRNAs (aa-tRNAs) and resulting in ribosome stalling. (**b**) Radezolid (green) also facilitates stalling of ribosomes with alanine in the penultimate position by enhancing radezolid binding to the PTC A-site. (**c**) Radezolid context-specificity is altered in Cfr-modified ribosomes due to repositioning of the C5 group required to accommodate the m^8^A2503 modification (pink circle). Rotation of the radezolid C5 group narrows the antibiotic pocket that normally houses the alanine side chain. Retained interaction between radezolid D-ring and A2602 (grey) explains why RZD can overcome Cfr-mediated resistance.

Our structural investigations into RZD binding to a Cfr-modified ribosome revealed the importance of the D-ring in stabilizing ribosome engagement, providing unprecedented insight into how RZD retains efficacy against Cfr modified ribosomes. Surprisingly, we found that while RZD retains inhibitory activity against Cfr-modified ribosomes, the m^8^A2503 modification disrupts RZD stalling on the tested alanine-containing peptide. Our structural findings suggest the Cfr modification likely changes the context specificity of RZD by remodeling the hydrophobic pocket engaged by alanine in complexes with *wildtype* ribosomes. In summary, our findings provide a unifying model for context-specific inhibition of translation by oxazolidinone antibiotics. Our observation that the antibiotic binding pocket is formed, in part, by the nascent peptide, has revealed the “missing” component of translation machinery involved in antibiotic binding from previous cross-linking experiments. The importance of Ala in the penultimate position of the nascent peptide for stabilizing the antibiotic provides the structural basis for context specificity observed in ribosome profiling and single molecule studies. Our structural insights into nascent-peptide specific inhibition of translation by oxazolidinones, and complementary work in related systems^19,37–39^, suggests prospects for the development of drugs that can modulate activity of the ribosomes in a protein-selective manner.

## DATA AVAILABILITY

Atomic coordinates for all presented structures have been deposited in the Protein Data Bank and EMDB under the following accession numbers:

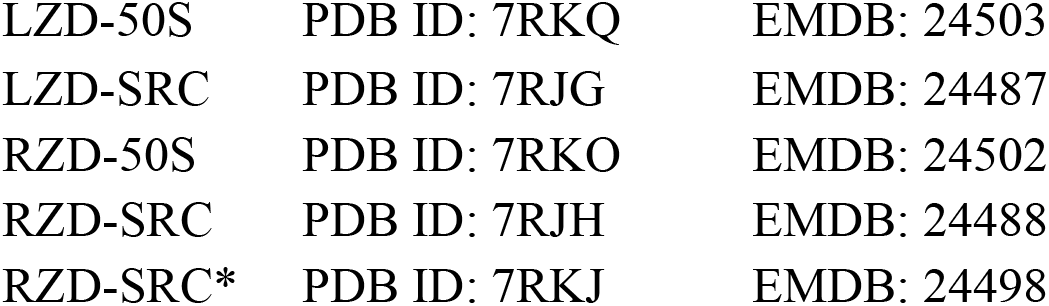

## ACKNOWLEDGEMENTS

We thank D. Bulkley and G. Gilbert for technical support at the UCSF Center for Advanced CryoEM, which is supported by the National Institutes of Health (S10OD020054 and S10OD021741) and the Howard Hughes Medical Institute (HHMI). Some of this work was performed at the Stanford-SLAC Cryo-EM Center (S^2^C^2^), which is supported by the National Institutes of Health Common Fund Transformative High-Resolution Cryo-Electron Microscopy program (U24 GM129541). The content is solely the responsibility of the authors and does not necessarily represent the official views of the National Institutes of Health. We thank E. Eng, E. Kopylov, and the rest of the staff for technical support at the National Center for CryoEM Access and Training (NCCAT) and the Simons Electron Microscopy Center located at the New York Structural Biology Center, which is supported by the NIH Common Fund Transformative High Resolution Cryo-Electron Microscopy program (U24 GM129539) and by grants from the Simons Foundation (SF349247) and NY State. We acknowledge support from NIAID (R01AI137270 to D.G.F and F32AI148120 to D.J.L.), W.M. Keck Foundation Medical Research Grant (to J.S.F. and D.G.F.), a Sanghvi-Agarwal Innovation Award (J.S.F.), NSF GRFP (1650113 to K.T.), and the UCSF Discovery Fellowship (to K.T).

## AUTHOR CONTRIBUTIONS

K.T. performed structural analysis, assisted with model refinement, prepared figures, and wrote the manuscript. V.S. prepared ribosome samples, performed model refinement, and edited the manuscript. D.J.L. performed cryo-EM analysis, performed model refinement, prepared figures, and edited the manuscript. I.D.Y. performed structural analysis, prepared figures, and assisted with model refinement. T.S. performed *in vitro* translation experiments. N.V.L. and A.M. provided data interpretation and edited the manuscript. J.S.F. and D.G.F. conceived and supervised the research, assisted in data interpretation, and edited the manuscript.

## METHODS

### Generation of DNA templates for PURExpress system

PCR reactions were conducted with AccuPrime Taq DNA Polymerase (Thermo Fisher). DNA templates used for toeprinting analysis were prepared as described previously^11^. DNA templates used for the generation of stalled ribosome complexes were prepared using by combining the following primers: 100 µM T7, 100 µM ORF_SD, 10 µM T7_MFKAF_Fwd, and 10 µM SD_MFKAF_Rev. Primer sequences are listed in **Supplementary Table 2**. The PCR product was purified using the MiniElute PCR kit (Qiagen) following manufacturer’s instructions and quality was confirmed using a 8% TBE (Novagen) gel. Sequence architecture of the resulting DNA product is outlined in **Supplementary Fig. 2**.

### Purification of 70S ribosomes

*Wildtype E. coli* 70S ribosomes were purified from the MRE600 strain as described previously^40^. Cfr-modified 70S ribosomes were prepared using previously published protocol with modification^33^ . In short, *E. coli* BW25113 expressing the evolved variant CfrV7 were grown to an OD600 of ∼0.7 in LB media containing ampicillin (100 µg/mL) and AHT inducer (30 ng/mL) at 37°C. After lysis using a microfluidizer, clarified lysates were applied to a 32% w/v sucrose cushion. Tight-coupled 70S ribosomes were purified on a 15-30% w/v sucrose gradient. To eliminate sucrose, ribosomes were precipitated by the addition of PEG 20,000 and subsequently resuspended in buffer containing 50 mM Hepes-KOH (pH 7.5), 150 mM KOAc, 20 mM Mg(OAc)_2_, 7 mM β-mercaptoethanol, 20 U/mL SuperASE-In.

### *In vitro* toeprinting assay

Toeprinting assays were conducted as previously described^11,38,41^. Briefly, reactions were prepared using the PURExpress **Δ** Ribosome Kit (New England Biolabs) in volumes of 5 µL and were allowed to proceed for 15 min at 37 °C. Primer extension was initiated by the addition of AMV reverse transcriptase (New England Biolabs) and allowed to proceed for 10 min. All reactions contained 50 µM mupirocin.

### Preparation of linezolid and radezolid-only bound ribosomes for cryo-EM

Purified 70S *wildtype* or Cfr-modified ribosomes were diluted to 1 pmol/µL in buffer containing 50 mM Hepes-KOH (pH 7.5), 150 mM KOAc, 20 mM Mg(OAc)_2_, 7 mM β-mercaptoethanol, 20 U/mL SuperASE-In. Diluted ribosomes were then incubated at 4°C for 1.5 h after the addition of either linezolid (Med Chem Express) or radezolid (Med Chem Express) in 40-60X molar excess. Samples were then filtered for 5 min at 14,000 x *g* at 4°C using a 0.22 uM low-binding Durapore PVDF filter (Millipore).

### Preparation of stalled ribosome complexes for cryo-EM

Stalled ribosome complexes containing the stalling peptide and corresponding oxazolidinone antibiotic were prepared by *in vitro* transcription-translation. Reactions of 100 µL volume were prepared using the PURExpress **Δ** Ribosome Kit (New England Biolabs, E3313S) containing 0.8 U/µL of SuperASE-In, ∼1100 ng of DNA template encoding the stalling peptide sequence (**Supplementary Fig. 2**), 5000 pmol of linezolid (Med Chem Express) or radezolid (Med Chem Express), and 360 pmol of *wildtype* ribosomes or 250 pmol of Cfr-modified ribosomes. The reaction was halted by placing reactions on ice after incubation at 37 °C for 1 h. The reaction was diluted to 190 µL by the addition of Buffer C (50 mM Hepes-KOH-pH 7.5, 150 mM KOAc, 20 mM Mg(OAc)_2_, 7 mM β-mercaptoethanol, 20 U/mL SuperASE-In) and purified by a 10-50% sucrose gradient also prepared in Buffer C. Ultracentrifugation was performed using a SW Ti41 rotor (Beckman Coulter) at 22,000 rpm for 16 h at 4°C. Gradients were fractionated using a Bio-Comp Fractionator in 20 fractions where absorbance at 260 nm was continuously monitored. Fractions corresponding to stalled ribosome complexes were precipitated by the slow addition of PEG 20,000 in Buffer C at 4°C to a final concentration of 8% w/v. Stalled complexes were isolated by centrifugation for 10 min at 17,500 x *g* at 4°C. After removing the supernatant, samples were slowly resuspended in Buffer C at 4°C. Sample concentration was determined by NanoDrop UV spectrophotometer (Thermo), where A_260_=1 corresponds to 24 pmol of 70S ribosome. Purified stalled ribosome complexes were then incubated with 20-30X molar excess of the corresponding oxazolidinone antibiotic (linezolid or radezolid) for 1 h at 4°C. Prior to freezing grids, stalled ribosome complexes were filtered for 5 min at 14,000 x *g* at 4°C using a 0.22 uM low-binding Durapore PVDF filter (Millipore).

### Cryo-EM analysis

Samples described above were diluted in Buffer C and deposited onto freshly glow-discharged (EMS-100 Glow Discharge System, Electron Microscopy Sciences, 30 s at 15 mA) copper Quantifoil (Quantifoil Micro Tools GmbH) grids with 2 nm thick amorphous carbon on top. Grids were incubated for 30 s at 10 °C and 95% humidity, before blotting and vitrification by plunging into liquid ethane using a FEI Vitrobot Mark IV (ThermoFisher). Ice thickness was controlled by varying the blot time, using Whatman #1 filter paper for blotting. Grids were screened for ice quality using a FEI Talos Arctica electron microscope (ThermoFisher, 200 kV, at UCSF) before grids were transported via dry shipper to other facilities or loaded into a UCSF FEI Titan Krios (ThermoFisher).

All datasets used for reconstruction were imaged on FEI Titan Krios microscopes (ThermoFisher, 300 kV). The LZD-50S and LZD-SRC datasets were collected at the Stanford-SLAC CryoEM Center (S^2^C^2^) using SerialEM on a microscope equipped with a Gatan K3 direct electron detector (DED) but without an imaging filter. The RZD-50S and RZD-SRC datasets were collected at the National Center for CryoEM Access and Training (NCCAT) using Leginon/Appion on a microscope equipped with a Gatan K2 Summit DED and an imaging filter (20 eV slit). The RZD-SRC* dataset was collected at UCSF on a microscope equipped with a Gatan K3 DED and an imaging filter (20 eV slit). The RZD-SRC* dataset was collected on-axis; all other datasets were collected using a nine-shot beam-image shift approach with coma compensation. All image stacks were collected in super-resolution mode. Pixel sizes, micrograph count, defocus values, and exposures varied slightly between facilities and are reported in **Supplementary Table 1**.

All image stacks were binned by a factor of 2, motion corrected, and dose-weighted using UCSF MotionCor2^42^. All reconstructions used dose-weighted micrographs. Initial CTF parameters were determined using CTFFIND4 within the cisTEM (v1.0.0-beta)^16^ software sweet. Micrographs with poor CTF fits or crystalline ice were excluded. Unsupervised particle picking used a soft-edged disk template was followed by 2D classification in cisTEM. Initial and final particle counts are reported in **Supplementary Table 1**. Only classes that clearly contained ice were omitted. An *ab initio* reconstruction was carried out in cisTEM on the RZD-SRC* dataset, which yielded a starting reference which was lowpass filtered and used as the initial reference for all five datasets. For SRC datasets, multi-class Auto refinement in cisTEM was used to select for all particles that had tRNA present. After this, and all non-SRC datasets, were subjected to a two-class Auto refinement in cisTEM to classify between “good” particles and damaged “garbage” particles and high-frequency noise. The good classes were carried forward into single class Auto and manual refinement efforts, including per-particle CTF estimation. Care was taken not to increment the high-resolution cutoff in refinement to prevent overfitting. Unsharpened maps were used in model refinement and figure preparation. Pixel size was confirmed by comparison and cross-correlation between the resulting map and a crystallographically-derived ribosome structure. 70S maps were used for model building, but 50S focused refinement (using a binary mask to select for the 50S in cisTEM) was carried out to aid in figure preparation. Map resolution values are reported as particle Fourier Shell Correlation (FSC) at 0.143.

### Atomic model building and refinement

Atomic models of 50S ribosomal subunit with antibiotics and 70S stalled ribosome complexes were generated by rounds of model building in Coot^43^ and refinement in PHENIX^44^. The atomic models of the *wildtype E. coli* 50S subunit (PDB 6PJ6) was used as the starting point. Initial models for the three 70S stalled ribosome complexes were obtained by combining: (i) a model of *wildtype E. coli* 50S subunit (PDB 6PJ6); (ii) a model of the 30S subunit from *wildtype E. coli* ErmBL-stalled ribosome structure (PDB: 5JU8)^45^; (iii) P-tRNA and mRNA extracted from the ErmBL-stalled ribosome structure (PDB: 5JU8, mutated and remodeled in *Coot*^*43*^ to yield fully modified *E. coli* tRNA^Phe^ and a short mRNA, respectively); and (iv) the nascent peptide, which was modelled in *Coot*. Model refinement against the acquired cryo-EM map was performed by multiple rounds of manual model building and restrained parameter-refinement (real-space refinement, positional refinement, and simulated annealing). Modified nucleotides were generated using eLBOW^46^ within Phenix^40^. The L3, L10 and L31 proteins were not modelled in any of the cryo-EM structures. Prior to running MolProbity^47^ analysis, nucleotides 76-94, 998-1009, and 1020-1040 of 16S rRNA, nucleotides 1053–1107, 2100–2189 of 23S rRNA, and ribosomal proteins L9 and L11 were removed, due to their high degree of disorder. Overall, protein residues and rRNA nucleotides show well refined geometrical parameters (**Supplementary Table 1**). Figures were prepared using Pymol Molecular Graphics System Version 2.4.1 Schrödinger, LLC or UCSF ChimeraX Version 1.2.5^48^.

