## Supplemental Materials for "Structural basis for context-specific inhibition of translation by oxazolidinone antibiotics"

***Short title:*** Translation inhibition by oxazolidinone antibiotics

**Authors:**

Kaitlyn Tsai[1]*, Vanja Stojković[1]*, D. John Lee[2]*, Iris D. Young[2], Teresa Szal[3,4], Nora Vazquez-Laslop[3,4], Alexander S. Mankin[3,4], James S. Fraser[2,5,#], Danica Galonić Fujimori[1,5,6,#]

[1] Department of Cellular and Molecular Pharmacology; University of California San Francisco, San Francisco, CA 94158, USA

[2] Department of Bioengineering and Therapeutic Sciences; University of California San Francisco, San Francisco, CA 94158, USA

[3] Department of Pharmaceutical Sciences, University of Illinois at Chicago, Chicago, IL 60607, USA

[4] Center for Biomolecular Sciences, University of Illinois at Chicago, Chicago, IL 60607, USA

[5] Quantitative Biosciences Institute, University of California San Francisco, San Francisco, CA 94158, USA

[6] Department of Pharmaceutical Chemistry, University of California San Francisco; San Francisco, CA 94158, USA

*Authors contributed equally to this work

#To whom correspondence should be addressed:

**This file includes:**

1. Supplementary Figs. 1-10
2. Supplementary Tables 1 & 2
3. Supplementary Information References

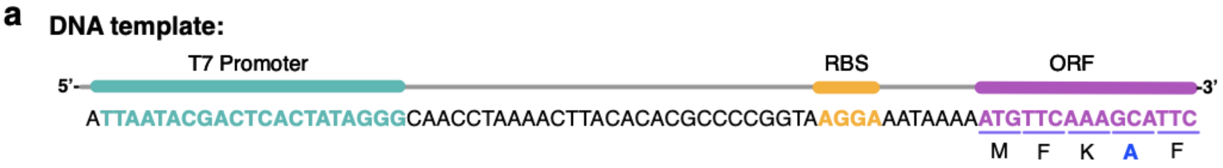


**Supplementary Fig. 1. Preparation of oxazolidinone-stalled ribosome complexes for cryo-EM analysis.** (**a**) Architecture of the DNA template encoding the T7 promoter, ribosome binding site (RBS), and model MFK**A**F stalling peptide open reading frame (ORF). Generation of stalled ribosomes at the F5 codon was biased by not including a stop codon at the end of the stalling peptide ORF.

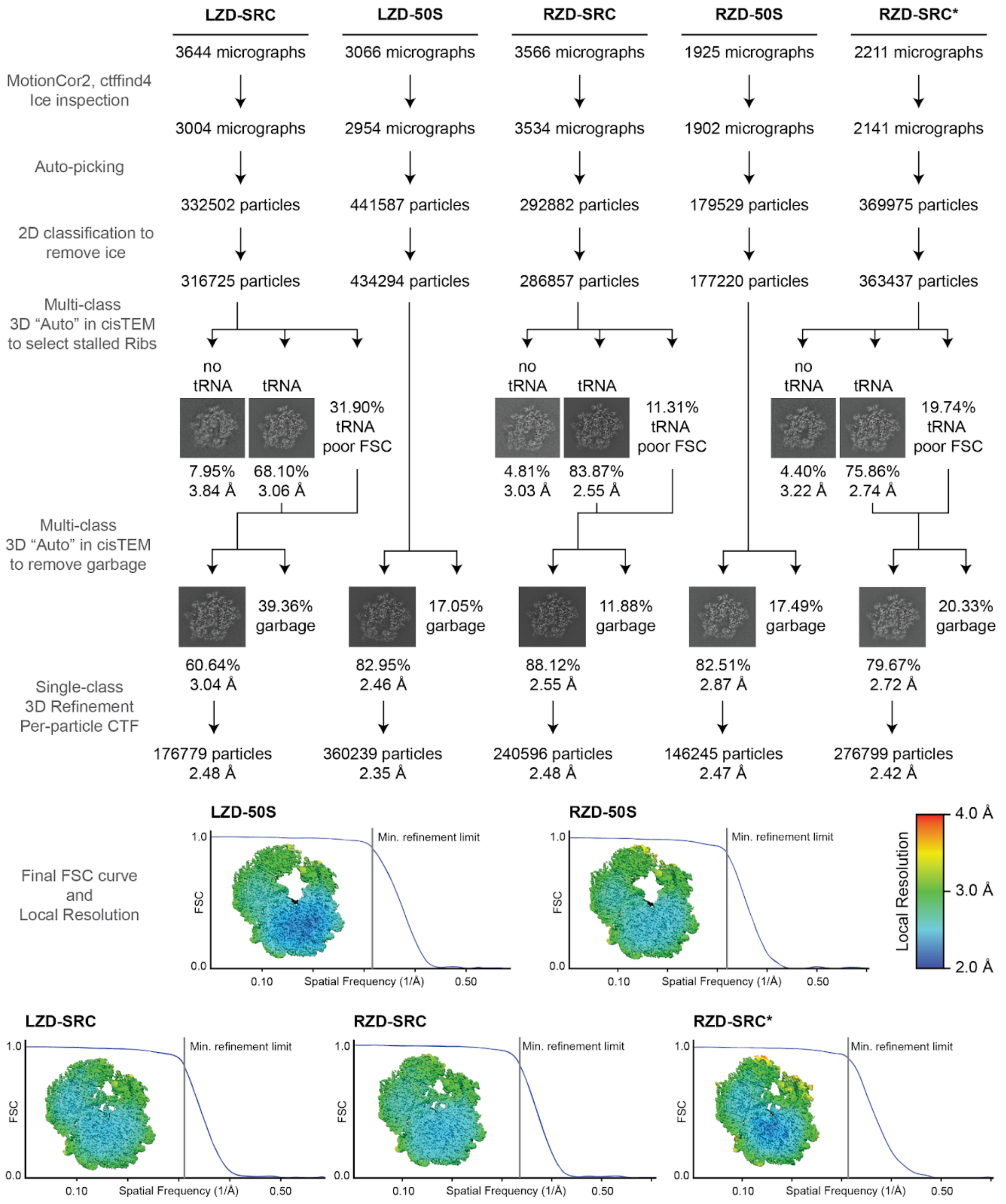


**Supplementary Fig. 2. Processing workflow, classification tree, FSC curves, and local resolution estimation for each map.** Micrographs were CTF corrected and curated for ice quality, followed by unsupervised particle picking. 2D classification was used to remove residual ice particles, with all others subjected to multi-class 3D classification and refinement approaches to remove particles without tRNA (for SRCs) and select for good particles. Final FSC curves are presented along with center-slab representation of local resolution. All steps were carried out within the cisTEM (v1.0.0-beta) framework with the exception of local resolution estimation, which used the Relion (v3.1.2) implementation.

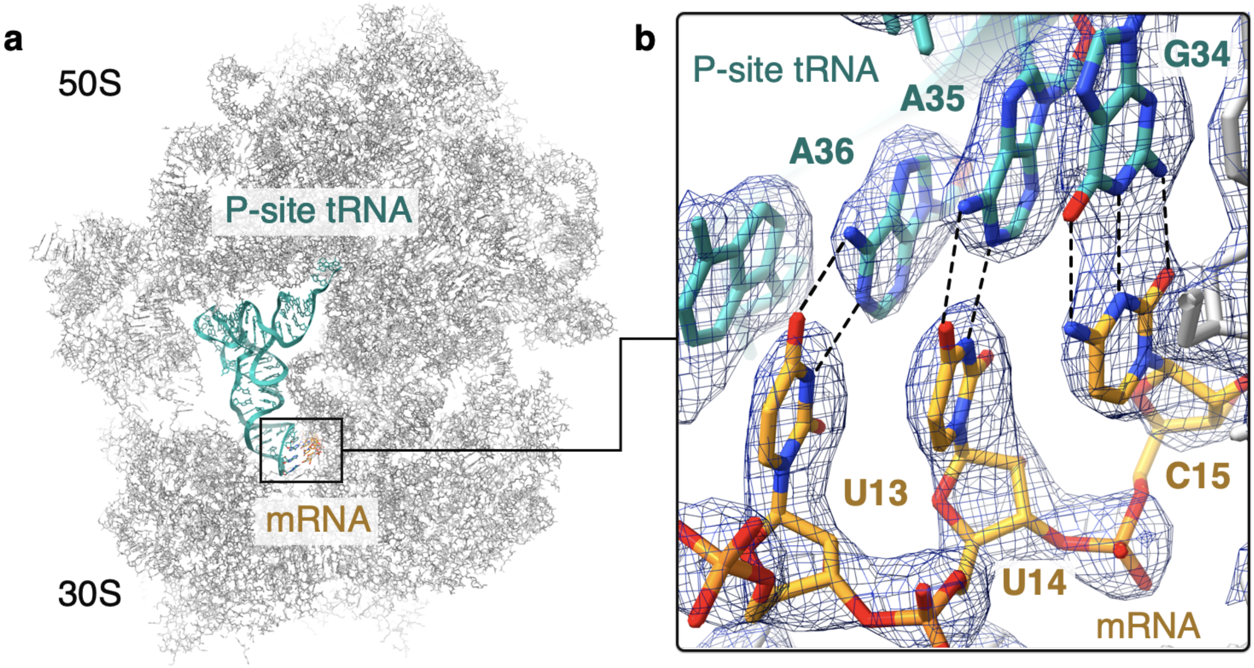


**Supplementary Fig. 3. Confirmation of oxazolidinone-induced ribosome stalling at the Phe5 codon.** (**a**) Location of the codon-anticodon interaction within the linezolid-stalled 70S ribosome complex. (**b**) Density for the codon-anticodon interaction is best modeled as UUC:GAA-tRNA$^{Phe}$ rather than GCA:UGC-tRNA$^{Ala}$ which would correspond to stalling at the upstream Ala4 codon. The figure was generated from unsharpened maps and the density is contoured at 4$\sigma$.

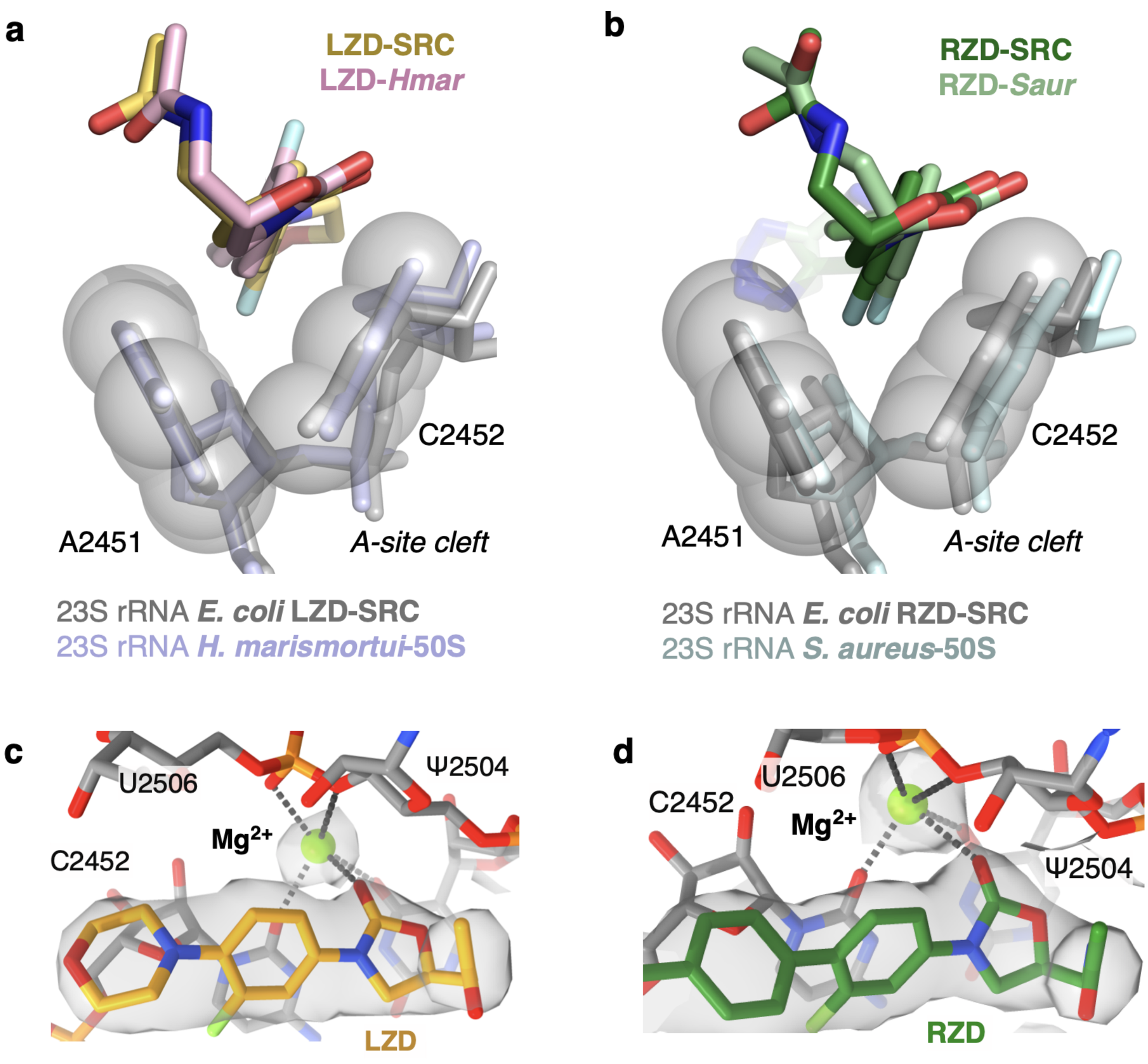


slack

**Supplementary Fig. 4. Overview of the binding mode of linezolid and radezolid within the ribosomal PTC.** (**a**) Structural overlay of linezolid (LZD) bound to a *H. marismortui* ribosome (PDB: 3CPW) and linezolid-stalled *E. coli* ribosome complex (LZD-SRC), highlighting binding within the A-site cleft. (**b**) Structural overlay of radezolid (RZD) bound to a *S. aureus* ribosome (PDB: 6WQQ) and radezolid-stalled *E. coli* ribosome complex (RZD-SRC), also highlighting the A-site cleft. Overlays for (**a**) and (**b**) were generated by aligning 23S rRNA nucleotides. (**c**) LZD oxazolidinone ring is coordinated to a $Mg^{2+}$ ion. Coulomb potential density is contoured at 2.5$\sigma$ from unsharpened density maps. (**d**) RZD oxazolidinone ring coordination to a $Mg^{2+}$ ion. Coulomb potential density is contoured at 3$\sigma$ from unsharpened density maps.

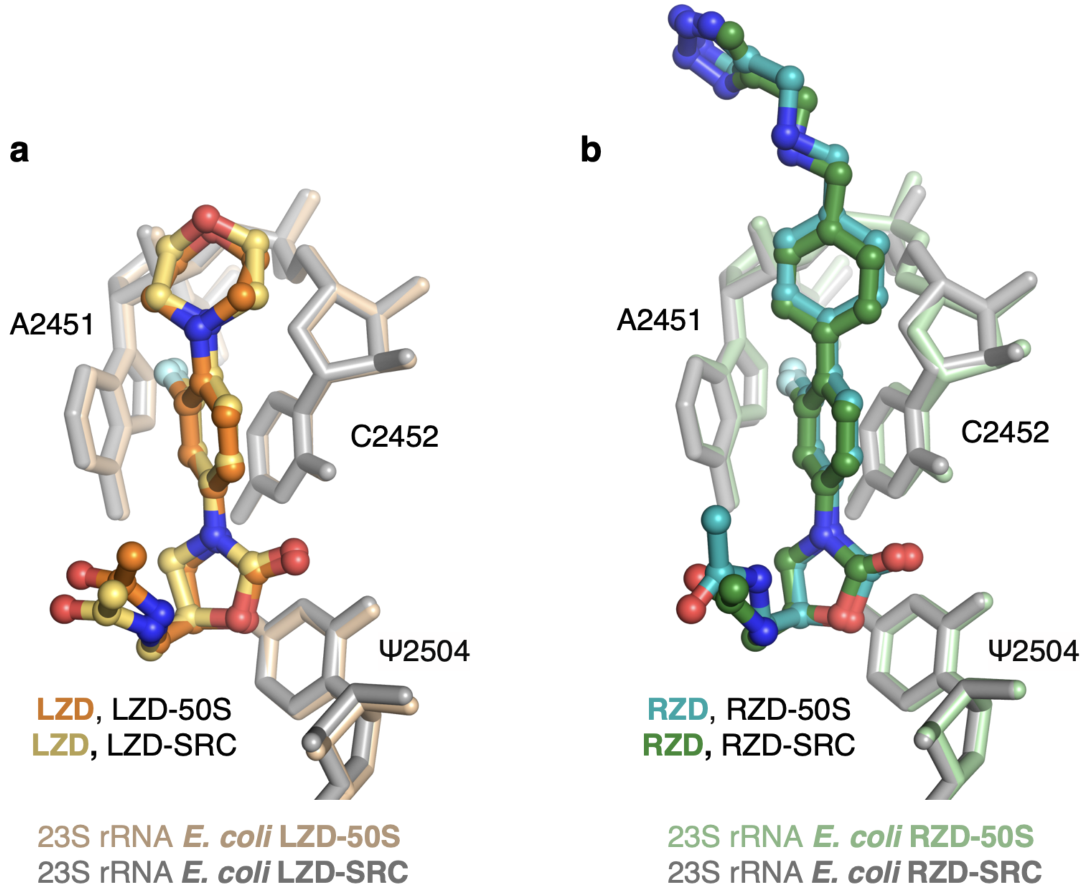


**Supplementary Fig. 5. Oxazolidinone binding modes in antibiotic-only and stalled ribosome complexes.** (**a**) Comparison of linezolid (LZD) binding modes in the antibiotic-alone (LZD-50S, antibiotic in orange) versus LZD-stalled complex (LZD-SRC, antibiotic in yellow). (**b**) Comparison of radezolid (RZD) binding modes in the antibiotic-alone (RZD-50S, antibiotic in teal) versus RZD-stalled complex (RZD-SRC, antibiotic in green). Structural overlays in this figure were performed by aligning the 23S rRNA chain.

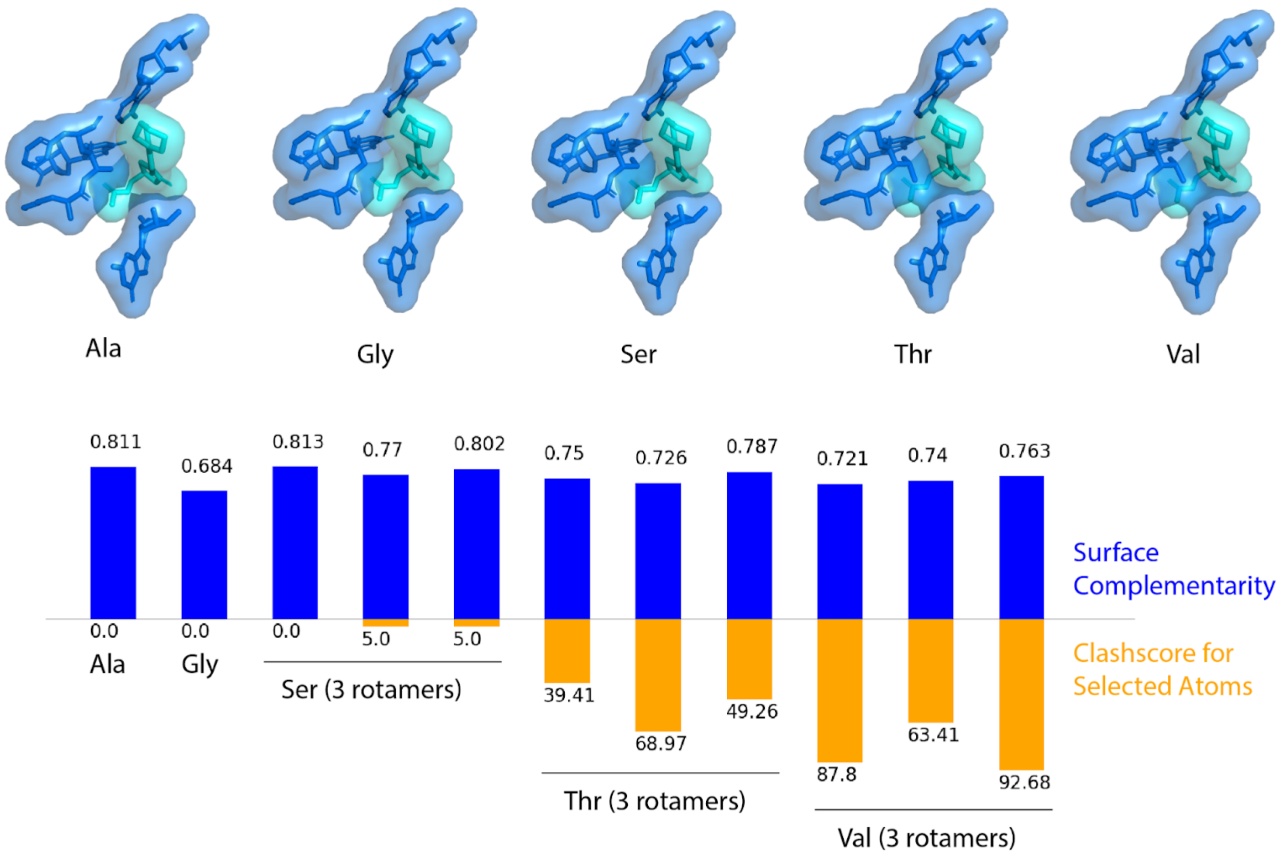


**Supplementary Fig. 6. *In silico* analysis of residues at the penultimate position.** Substitution of glycine (Gly), serine (Ser), threonine (Thr), or valine (Val) for alanine (Ala) at the penultimate position results in reduced surface complementarity $S_c$ as measured by the sc tool in CCP4[1] for all except one rotamer of serine, which produces a slightly higher score. Surface complementarity is calculated between two selections of atoms, the first of which is the sequence of three residues centered on the penultimate position, and the second of which is the linezolid ligand and the three nucleotides close enough to potentially interact either favorably or disfavorably with the peptide sequence in the first selection. Alanine and one rotamer of serine are most highly favored for ligand binding by this metric. Clashscore is also calculated on the set of all atoms in both selections. Substitutions larger than serine produce clashscores indicative of occlusion of the ligand.

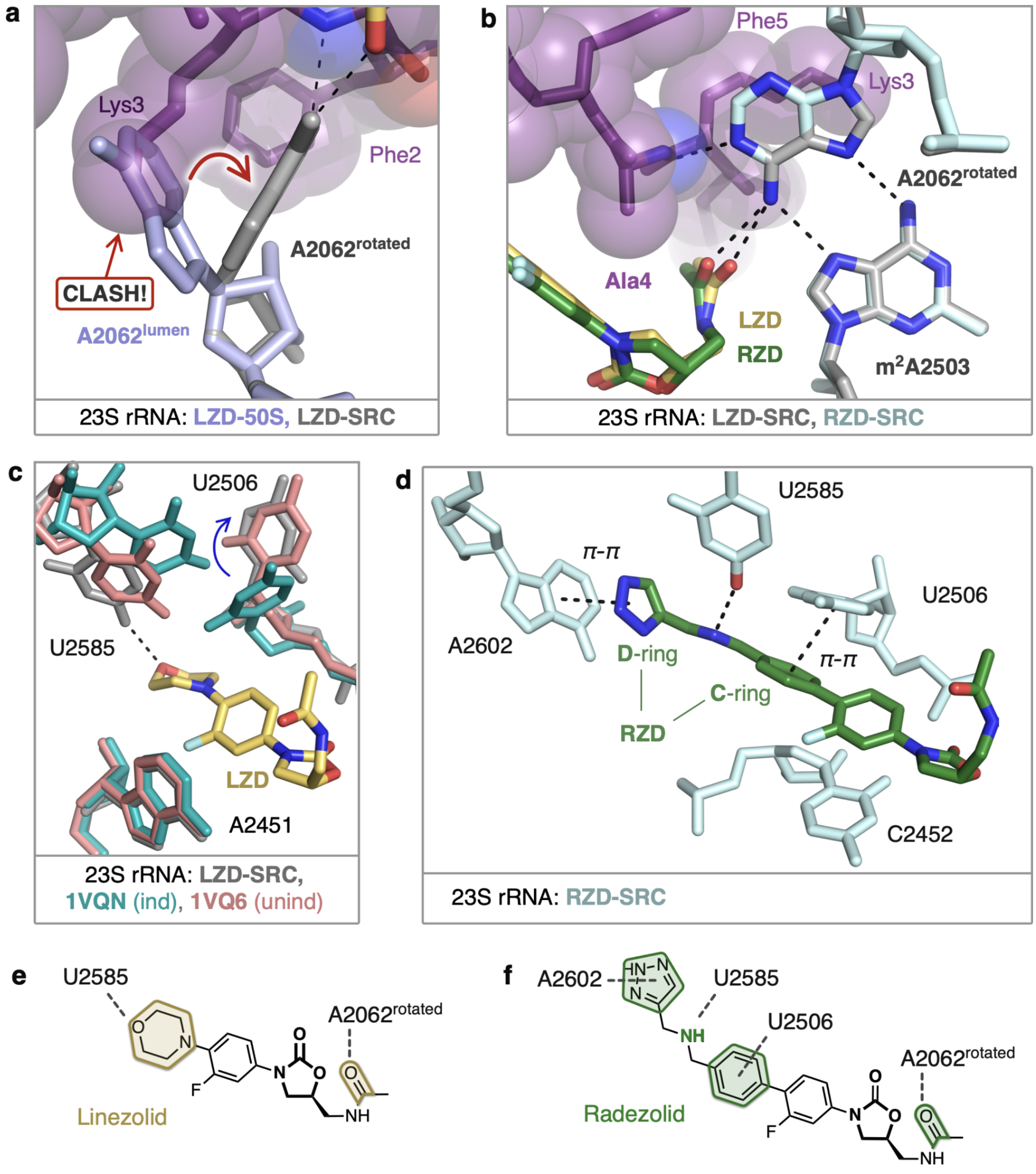


**Supplementary Fig 7. In the stalled complexes, rRNA nucleotides provide additional contacts with oxazolidinone antibiotics.** (**a**) The rotated conformation of A2062 is adopted to accommodate the nascent chain (purple). (**b**) In the rotated state, A2062 makes H-bond interactions with the nascent chain (purple), m$^2$A2503, and the C5 group of oxazolidinone antibiotic linezolid (LZD, yellow) and radezolid (RZD, green). (**c**) Additional interaction between U2585 and the morpholine ring of LZD (yellow). U2506 adopts a conformation consistent with the unaccommodated (unind, pink) state of the PTC[2]. The accommodated PTC state (ind, teal) shown for comparison[2]. (**d**) Stabilized interactions between RZD and A2602, U2585, and U2506 observed in the stalled complex. (**e,f**) Schematics of stabilized interactions for (panel e) linezolid and (panel f) radezolid.

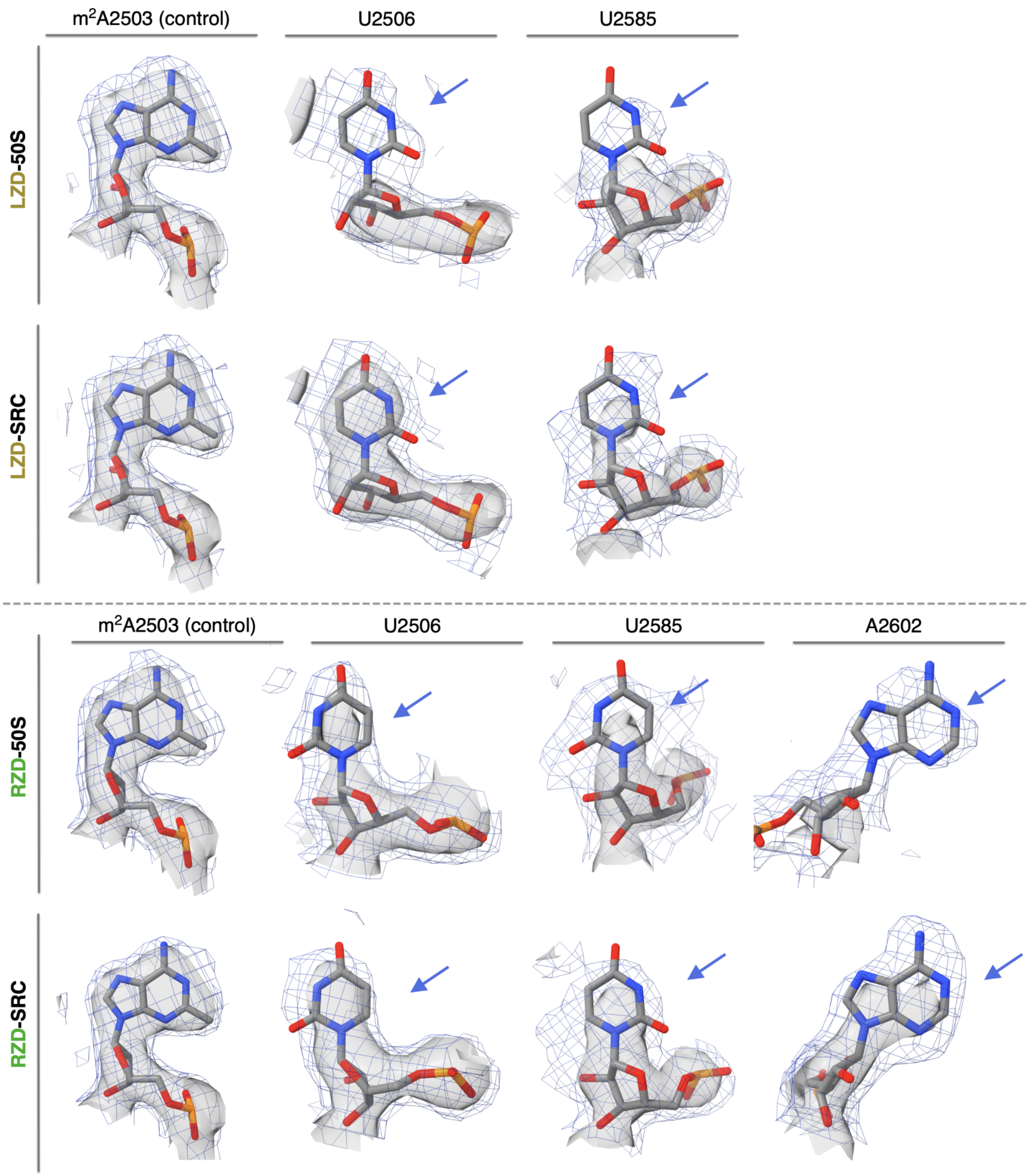


**Supplementary Fig. 8. Stabilization of PTC nucleotides in the oxazolidinone-stalled ribosome complexes.** Direct comparison of nucleotide density in the linezolid-only bound (LZD-50S) and the linezolid-stalled complex (LZD-SRC), displaying $m^2$A2503 as a control nucleotide and dynamic nucleotides U2585 and U2506. Direct comparison of nucleotide density in the radezolid-only bound (RZD-50S) and the radezolid-stalled complex (RZD-SRC), displaying A2503 and dynamic nucleotides U2506, U2585, A2602. Coulomb potential density is contoured at 4.0$\sigma$ in surface representation and 1.0$\sigma$ in mesh representation from unsharpened cryo-EM density maps generated with a soft 50S mask.

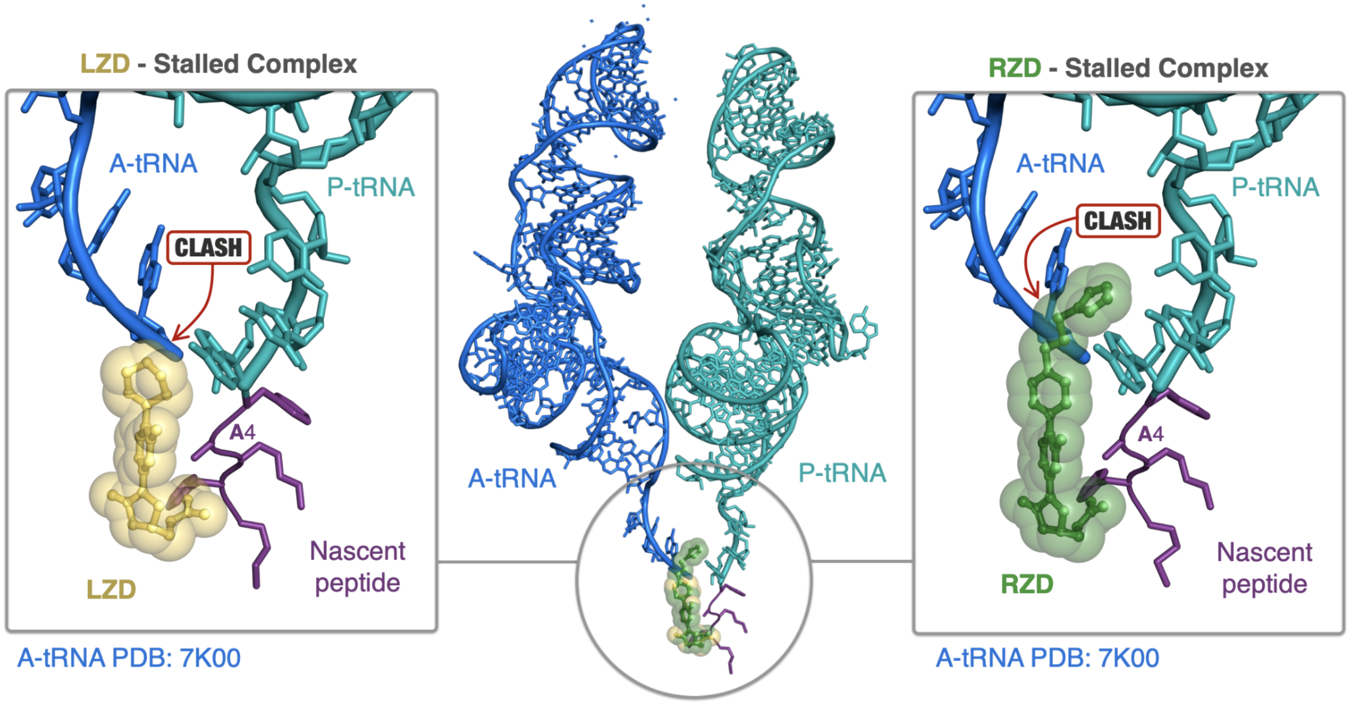


**Supplementary Fig. 9. Steric occlusion of A-site tRNA binding by oxazolidinone antibiotics.** Linezolid (LZD, yellow) and radezolid (RZD, green) prevent binding of A-site tRNAs (blue, from PDB: 7K00). Prominent steric clashes between the A-tRNA and LZD C-ring and RZD C/D-ring are highlighted. Structural overlays were performed by alignment of 23S rRNA nucleotides.

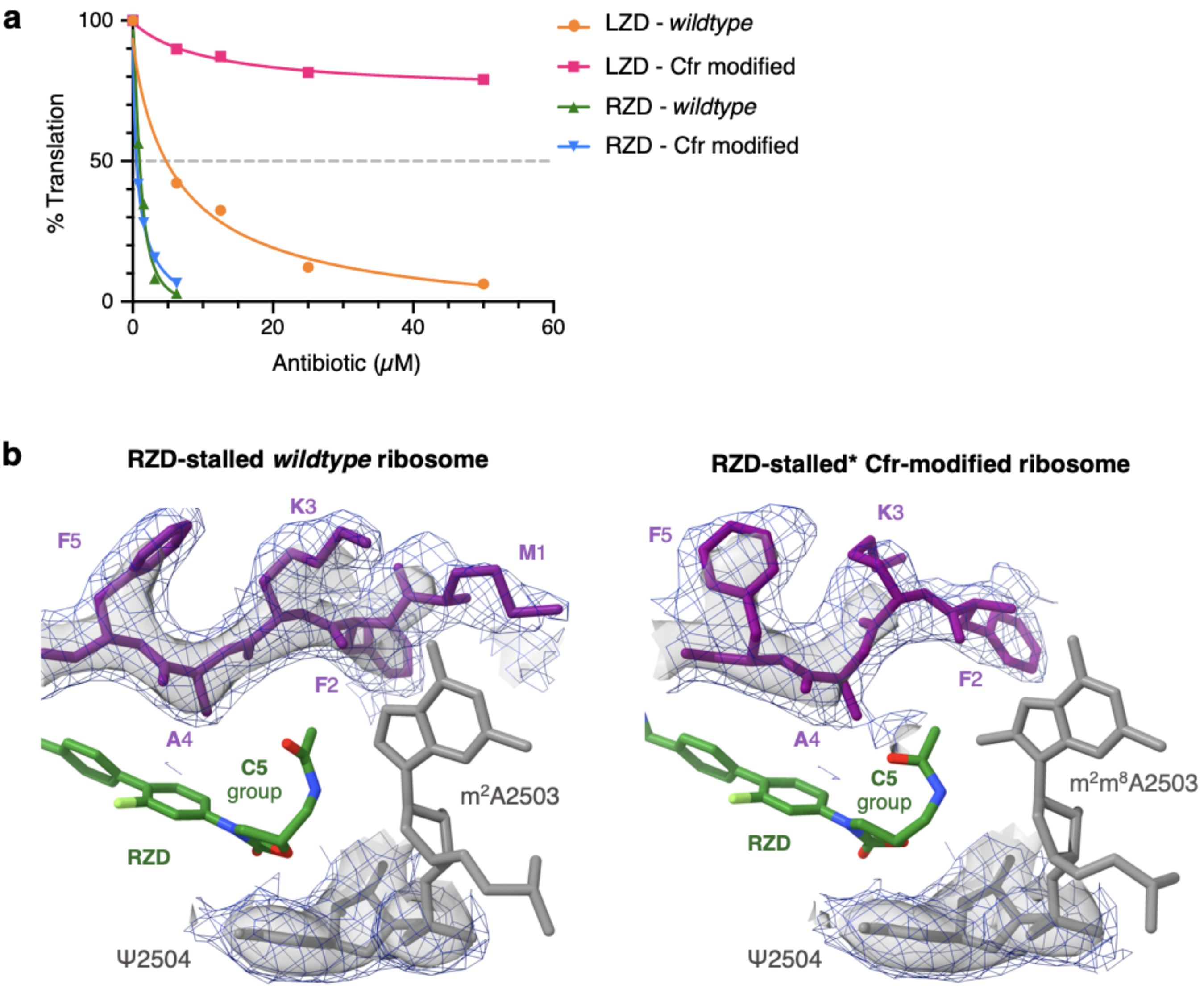


**Supplementary Fig. 10. RZD inhibitory activity and binding mode with Cfr-modified ribosomes.** (**a**) *In vitro* activity of linezolid (LZD) and radezolid (RZD) against *wildtype* and Cfr-modified ribosomes determined by inhibition of sfGFP translation. Percent (%) translation calculated as percentage of sfGFP translation at the tested antibiotic concentration compared to reactions containing no antibiotic. (**b**) Density comparison for the alanine-containing MFK**A**F stalling peptide (purple) between RZD-stalled complexes with *wildtype* (left) and Cfr-modified (right) ribosomes. RZD shown in green and 23S rRNA nucleotides shown in grey. Density for Ψ2504 shown as a control. Coulomb potential density is contoured at 4.0$\sigma$ in surface representation and 1.0$\sigma$ in mesh representation from unsharpened maps generated with a soft 50S mask. Of note, the side chain for Met1 of the nascent chain was not modeled in the Cfr-modified ribosome (right).

| **Structure Name**<br>PDB<br>EMDB | **LZD-SRC**<br>PDB: 7RJG<br>EMDB: 24487 | **LZD-50S**<br>PDB:7RKQ<br>EMDB: 24503 | **RZD-SRC**<br>PDB: 7RJH<br>EMDB: 24488 | **RZD-50S**<br>PDB: 7RKO<br>EMDB: 24502 | **RZD-SRC***<br>PDB: 7RKJ<br>EMDB: 24498 |
|---|---|---|---|---|---|
| **Data collection and processing** | | | | | |
| Facility and Electron microscope | $S^2C^2$ Titan Krios | $S^2C^2$ Titan Krios | NCCAT Titan Krios | NCCAT Titan Krios | UCSF Titan Krios |
| Voltage (kV) | 300 | 300 | 300 | 300 | 300 |
| Camera | Gatan K3 | Gatan K3 | Gatan K2 Summit | Gatan K2 Summit | Gatan K3 |
| Nominal Magnification | 29,000 | 29,000 | 105,000 | 105,000 | 105,000 |
| Electron dose (e-/Å$^2$) | 67.8 | 67.8 | 52.8 | 52.8 | 69.0 |
| Defocus range (μm) | 0.3-0.8 | 0.3-0.8 | 0.5-1.5 | 0.5-1.5 | 0.3-0.8 |
| Pixel size (Å) | 0.8125 | 0.8125 | 0.8250 | 0.8250 | 0.8261 |
| Energy filter slit width (eV) | n/a | n/a | 20 | 20 | 20 |
| Symmetry imposed | C1 | C1 | C1 | C1 | C1 |
| Number of total micrographs | 3644 | 3066 | 3566 | 1925 | 2211 |
| Number of good micrographs | 3004 | 2954 | 3534 | 1902 | 2141 |
| Number of particles picked from good micrographs | 332502 | 441587 | 292882 | 179529 | 369975 |
| Number of particles used in final reconstruction | 176779 | 360239 | 240596 | 146245 | 276799 |
| **Map refinement** | | | | | |
| Model resolution (Å) | 2.48 | 2.35 | 2.48 | 2.47 | 2.42 |
| FSC threshold | 0.143 | 0.143 | 0.143 | 0.143 | 0.143 |
| Map sharpening B-factor (Å$^2$) | 0.0 | 0.0 | 0.0 | 0.0 | 0.0 |
| **Refinement and model statistics** | | | | | |
| Model composition | | | | | |
| Total atoms | 241368 | 151158 | 205313 | 151126 | 205269 |
| Hydrogens only | 96909 | 60046 | 60872 | 60028 | 60862 |
| Non-hydrogen atoms | 144459 | 91112 | 144441 | 91098 | 144407 |
| Protein residues | 5715 | 3356 | 5715 | 3356 | 5714 |
| Nucleotide residues | 4637 | 3016 | 4637 | 3015 | 4636 |
| Ligands | 182 | 183 | 185 | 182 | 172 |
| Clashscore, all atoms | 7.24 | 5.76 | 7.7 | 6.2 | 5.8 |
| Protein geometry | | | | | |
| MolProbity score | 2.15 | 1.93 | 1.94 | 2.04 | 1.83 |
| Rotamer outliers (%) | 2.68 | 2.49 | 1.15 | 3.09 | 1.10 |
| Cβ deviations >0.25 Å (%) | 0 | 0.04 | 0.02 | 0 | 0.04 |
| Ramachandran (%) | | | | | |
| - Favored | 93.41 | 95.52 | 92.25 | 95.32 | 92.25 |
| - Allowed | 6.1 | 4.44 | 6.29 | 4.65 | 6.29 |
| - Outliers | 0.49 | 0.03 | 1.46 | 0.03 | 1.46 |
| Deviations from ideal geometry | | | | | |
| - Bonds (%) | 0 | 0.01 | 0.39 | 0 | 0.01 |
| - Angles (%) | 0.01 | 0.03 | 0.14 | 0.02 | 0.01 |
| Nucleic acid geometry | | | | | |
| Probably wrong sugar puckers (%) | 1.06 | 0.94 | 1.13 | 1.15 | 1.26 |
| Bad backbone conformations (%) | 16.37 | 15.19 | 18.71 | 17 | 18.75 |
| Bad bonds (%) | 0.03 | 0.02 | 0.39 | 0.01 | 0.40 |
| Bad angles (%) | 0.11 | 0.00 | 0.14 | 0 | 0.17 |
| EM Ringer Score | 3.0 | 3.4 | 2.0 | 3.2 | 1.7 |

**Supplementary Table 1. Cryo-EM data collection and statistics.** Structural information for the following *E.coli* ribosome structures: linezolid-stalled ribosome complex (LZD-SRC), linezolid-only bound (LZD-50S), radezolid-stalled ribosome complex (RZD-SRC), radezolid-only bound (RZD-50S), and radezolid-artificially stalled on Cfr-modified ribosomes (RZD-SRC*). Model composition is reported for full deposited models, molprobity metrics are for the trimmed models as reported in the methods section.

| Primer name | Sequence |
|---|---|
| T7 | 5’-ATTAATACGACTCACTATAGG-3’ |
| ORF_SD | 5’-GAATGCTTTGAACATTTTTATTTCC-3’ |
| T7_MFKAF_ Fwd | 5’-ATTAATACGACTCACTATAGGGCAACCTAAAACTTACACACGCCCCG-3’ |
| SD_MFKAF_Rev | 5’-GAATGCTTTGAACATTTTTATTTCCTTACCGGGGCGTGTGTAAGTTTTAG-3’ |

**Supplementary Table 2. Primer sequences used in this study.** Oligonucleotides were purchased from Integrated DNA Technologies and prepared with standard desalting procedures.

### SUPPLEMENTARY INFORMATION REFERENCES

1. Lawrence, M. C. S C: measuring shape complementarity at protein-protein interfaces. *CCP4 NEWSLETTER ON PROTEIN CRYSTALLOGRAPHY*.
2. Schmeing, T. M., Huang, K. S., Strobel, S. A. & Steitz, T. A. An induced-fit mechanism to promote peptide bond formation and exclude hydrolysis of peptidyl-tRNA. *Nature* **438**, 520–524 (2005).
